# The Role of Temperature on the Development of Circadian Rhythms in Honey Bee Workers

**DOI:** 10.1101/2020.08.17.254557

**Authors:** Manuel A. Giannoni-Guzmán, Janpierre Aleman-Rios, Alexander M. Melendez Moreno, Gabriel Diaz Hernandez, Melina Perez, Darimar Loubriel, Tugrul Giray, Jose L. Agosto-Rivera

**Affiliations:** Department of Biological Sciences, Vanderbilt University, Nashville, TN, USA; Department of Biology, University of Puerto Rico Rio Piedras Campus, San Juan, PR

**Keywords:** Honey bee, development, circadian, workers, temperature

## Abstract

Circadian rhythms in honey bees are involved in various processes that impact colony survival. For example, young nurses take care of the brood constantly throughout the day and lack circadian rhythms, while foragers use the circadian clock to remember and predict food availability in subsequent days. Previous studies suggested that development of circadian rhythms both in the field and the laboratory began around 7-9 days of age. However, not much is understood about the postembryonic development of circadian rhythms in honey bees. In the current study, we examine the effects of socially regulated colony temperature on the ontogeny of circadian rhythms of young workers under controlled laboratory conditions. We hypothesized that temperature plays a key role in the development of circadian rhythmicity in young workers. Our results show that young workers kept at 35°C develop circadian rhythmicity faster and in greater proportion than bees kept at 25°C. In addition, we examine if the effect of colony temperature during the first 48 hours after emergence is enough to observe effects on the rate and proportion of development of circadian rhythmicity. We observed that twice as many individuals that were exposed to 35°C during the first 48 hours develop circadian rhythms compared to individuals kept at 25°C. In addition, we observed differences in the average endogenous period length consistent with temperature compensation of the circadian rhythms between the 25°C and 35°C cohorts. We also observed differences in the degree of period length variation between the 25°C and 35°C cohorts, which combined with the proportion of arrhythmic individuals and survival data suggest that development of circadian rhythms is incomplete in individuals exposed to 25°C adult emergence. This study shows that temperature, which is socially regulated inside the hive, is a key factor that influences the ontogeny of circadian rhythmicity of workers.

## Introduction

The circadian clock of honey bees is important in complex physiological processes, such as spatiotemporal learning, time perception and sun-compass navigation (Goodwin and Lewis, 1987; Moore et al., 1998; Van Nest and Moore, 2012; von Frisch, 1967; Wagner et al., 2013). However, when it comes to development of circadian rhythms in honey bee workers, scientists are just beginning to scratch the surface of what is thought to be a highly complex mechanism of regulation, with components at the environmental, social, hormonal and genetic levels (Eban-Rothschild et al., 2012; Moore, 2001; Moore et al., 1998; Shemesh et al., 2007). In this manuscript, we study the role of environmental temperature on the ontogeny of circadian rhythms of young honey bee workers.

The development of honey bee circadian rhythms is of particular interest because similar to human infants, young honey bees present postembryonic development of circadian rhythms before they forage (Eban-Rothschild et al., 2012; Moore et al., 1998). Furthermore, in the colony, it is thought that workers will remain arrhythmic performing in-hive tasks and will develop circadian rhythmicity just prior to the onset of foraging behavior, suggesting that ontogeny of circadian rhythms is intertwined with age-related division of labor in the colony. Studies examining the timing of in-hive tasks such as brood care found that individual ‘nurses’ performed this task around the clock, which is thought to benefit the developing brood (Moore et al., 1998).

In isolation, during the first days of their adult life young bees lack behavioral, metabolic or daily oscillations in circadian gene expression in the brain, that are associated with circadian rhythmicity. Under these constant conditions (DD, ~60%RH, 26-30°C), researchers have reported that ontogeny of circadian rhythmicity occurs at around 7-10 days of age in 50% of the sampled subjects (Moore, 2001; Toma et al., 2000). Furthermore, under these experimental conditions by 16 days of age around 25% of the bees were still arrhythmic. Since ontogeny of circadian rhythms is thought to be regulated by age-related division of labor, researchers have manipulated neuroendocrine signals known to accelerate onset of foraging (such as juvenile hormone, octopamine and cGMP dependent protein kinase), hypothesizing a similar effect on circadian rhythms without success in individually isolated bees (Ben-Shahar, 2003; Bloch et al., 2002; Meshi and Bloch, 2007). A recent study examined whether the colony environment or other social cues may elicit strong circadian rhythms in young workers (Eban-Rothschild et al., 2012). Their findings reveal that experiencing the colony environment, either in a mesh cage or interacting with other bees for 48 hours after adult emergence, resulted in strong circadian rhythms when bees were brought to the laboratory. The authors of this work postulate that social cues, the colony microenvironment or a combination of both plays a role in the ontogeny of circadian rhythms of young workers. Taken together, these studies suggest the existence of a cue, which can be social or environmental, that elicits the development of circadian rhythmicity.

Honey bee colonies are able to efficiently regulate the colony microenvironment (Jones et al., 2004, 2007; Kronenberg and Heller, 1982; Seeley, 1974; Simpson, 1961). Studies have shown that bees regulate CO_2_ levels, humidity and temperature inside the colony (Ohashi et al., 2008). In response to an increase in CO_2_ levels inside the colony honey bee workers begin fanning until CO_2_ levels diminish (Seeley, 1974). While the ability of honey bees to control temperature has been the main interest of researchers, humidity inside the nest is also regulated by workers (Human et al., 2006). Studies have shown that colonies with a naturally mated queen, are able to regulate temperature better than colonies that originate from a single drone artificially inseminated queen (Jones et al., 2004). This temperature control is especially important, since deviations of more that 1.5°C from 35°C at the core of the hive during larval and pupal development can have lasting changes in the adult honey bee (Winston, 1987).

Environmental temperature is also important for locomotor activity rhythms. Studies examining the endogenous rhythms of the Japanese honey bee *Apis cerana* show that environmental temperature has a direct effect on the endogenous period length of foragers (Fuchikawa and Shimizu, 2007). Recent work in our laboratory using the gentle Africanized honey bee (*gAHB*) also shows that environmental temperature affects the endogenous period length in honey bee foragers (Giannoni-Guzmán et al., 2014). However, the effect of temperature in the development of circadian rhythms in honey bee workers has yet to be explored.

In the current study we examined the effects of environmental temperature on the development of circadian rhythms in young workers. We hypothesized that temperature at the center of the colony is important for the development of circadian rhythms in young honey bee workers. In order to test this hypothesis, we isolated 1-day-old workers in locomotor activity monitors either at 25°C or 35°C. We examined the endogenous period length of rhythmic individuals in each group, the variation in period length and the mortality between the groups. Lastly, given the previous body of work that indicates that the first 48 hours after emergence are important for the development of circadian rhythms, we examined the effect of colony temperature during these 48 hours by placing individuals at 35°C and then changing the temperature to 25°C. Our results highlight the importance of socially regulated temperature of the hive in the ontogeny of circadian rhythms in honey bee workers.

## Materials and Methods

### Honey bees Colonies and collections

Colonies used in our experiments had mated queens that were laying eggs of gentle Africanized honey bees (Gallindo-Cardona et al., 2013). These colonies were located at the University of Puerto Rico (UPR) Gurabo Experimental Station in Gurabo, Puerto Rico. For all experiments, brood frames were collected, workers were removed and then the frame was stored in an incubator overnight (~35°C). The following morning, bees that emerged from the frames were collected and placed inside individual tubes for locomotor activity monitoring. The first colony of experiment 1 was examined on November 29, 2012 (colony 1), while the second colony was assayed beginning January 12, 2013 (colony 2). A total of 320 bees were used in this experiment, 256 for colony 1 and 64 for colony 2. Experiment 2 examined the effect of temperature during the first 48 hours after eclosion on the development of circadian rhythms, fixed began on February 26, 2016.

### Experiment 1: Development of Circadian rhythms at 25°C vs. 35°C

Locomotor activity measurements were carried out using two environmental chambers (Percival, I-30BLL) set up under constant darkness, relative humidity of 80%±5% and temperature of 25±0.5°C or 35±0.5°C and maintained constant throughout the experiments. Locomotor activity was recorded using monitors and software from Trikinetics (Waltham, MA, USA) as previously described (Giannoni-Guzmán et al., 2014). Briefly, 1-day-old workers were collected from the brood frame and placed inside individual tubes within the activity monitoring system. Food in the form of honey candy (mixed sugar and honey) and water were provided “*ad-libitum*” and changed as needed. Circadian rhythmicity was determined using 4 consecutive days of data (days 6-10), using autocorrelation analysis for 1-minute bins (Levine et al., 2002). All bees were approximately the same age for periods where rhythmicity was analyzed.

### Experiment 2: Development of circadian rhythms after 48 hours at 35°C

As in experiment 1, we carried out locomotor activity measurements using two environmental chambers. In one of these the temperature during the first 48 hours was set at 35°C and afterwards lowered to 25°C for the remainder of the experiment. The other incubator was kept at 25°C throughout the experiment. Food and water where provided *ad libitum* and changed as needed.

### Data analysis

All data sets were tested for normality via a Goodness of Fit test and appropriate nonparametric statistics were used were needed. The locomotor activity of each individual was processed using freely available MatLab^®^ toolboxes developed in Jeffrey Hall’s laboratory (Levine et al., 2002). Visual examination of locomotor activity for each individual in the form of actograms was utilized to determine the age at onset of circadian rhythms. Repeated measures MANOVA were utilized to determine if there were significant differences between the onset of rhythmicity between each of the experimental groups. Autocorrelation plots were utilized to confirm rhythmicity and calculate period length for each bee. Period length analysis was calculated for days. To examine differences in average period length between cohorts a two-way ANOVA was performed. To determine differences in the degree of period length variation the Levine’s test for equality of variance was performed.

To determine if environment temperature influences survival in our experiments, we performed survival analysis via the Gehan-Breslow-Wilcoxon test. Furthermore, Proportional Hazards analysis was performed to determine if differences in mortality were the result of independent factors or a combination of different factors. All statistical analyses were performed using the JMP™ software package from SAS (SAS Institute Inc., 2009); graphs and figures were created in MATLAB (MathWorks, Inc., Natick, MA, USA) and GraphPad Prism 6.00 (GraphPad Software, La Jolla, CA, USA).

## Results

Consistent with the hypothesis that brood nest temperature is important for the ontogeny of circadian rhythms, our results show that young workers kept at 35°C developed circadian rhythms as early as 2 days of age compared to young workers kept at 25°C, which began developing rhythms between 4-5 days of age (Figure 1). In addition, at 35°C between 60-80% of workers developed circadian rhythms, while at 25°C less than 30% of the bees developed rhythmicity (Repeated measures MANOVA, colony 1: F=3.94, df=9, p<<0.001; colony 2: F=3.29, df=7, p<0.01) (Figure 1). This result indicates that colony temperature plays a key role in the development of circadian rhythmicity. Further examination of locomotor activity plots of individuals that developed circadian rhythms revealed not only that the onset of circadian rhythmicity was different between groups, also that the endogenous period length in young workers was different between each experimental group (Figure 2).

**Figure 1.**
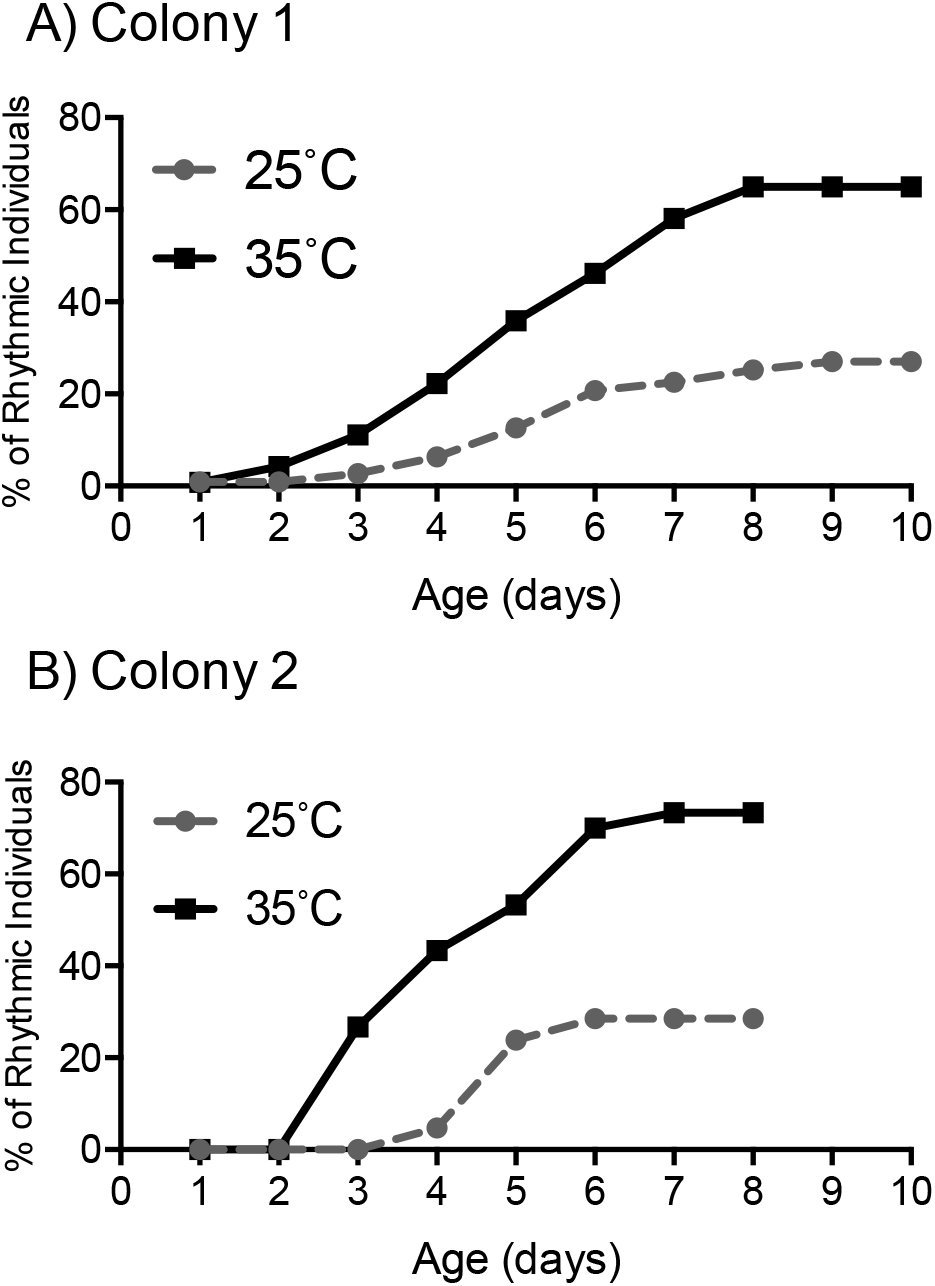
Rate and proportion of young workers developing circadian rhythms is greater at 35°C than at 25°C. Cumulative distribution of rhythmic young workers at 25°C and 35°C in constant darkness for two colonies. At 35°C the rate of development and the proportion of 1-day-old bees developing strong circadian rhythms were higher than at 25°C. Repeated measures MANOVA for each of the colonies samples yielded significant differences between the 25°C and 35°C conditions for both colonies sampled **A)** Colony 1 (F=3.94, df=9, p<<0.001). **B)** Colony 2 (F=3.29, df=7, p<0.01).

**Figure 2.**
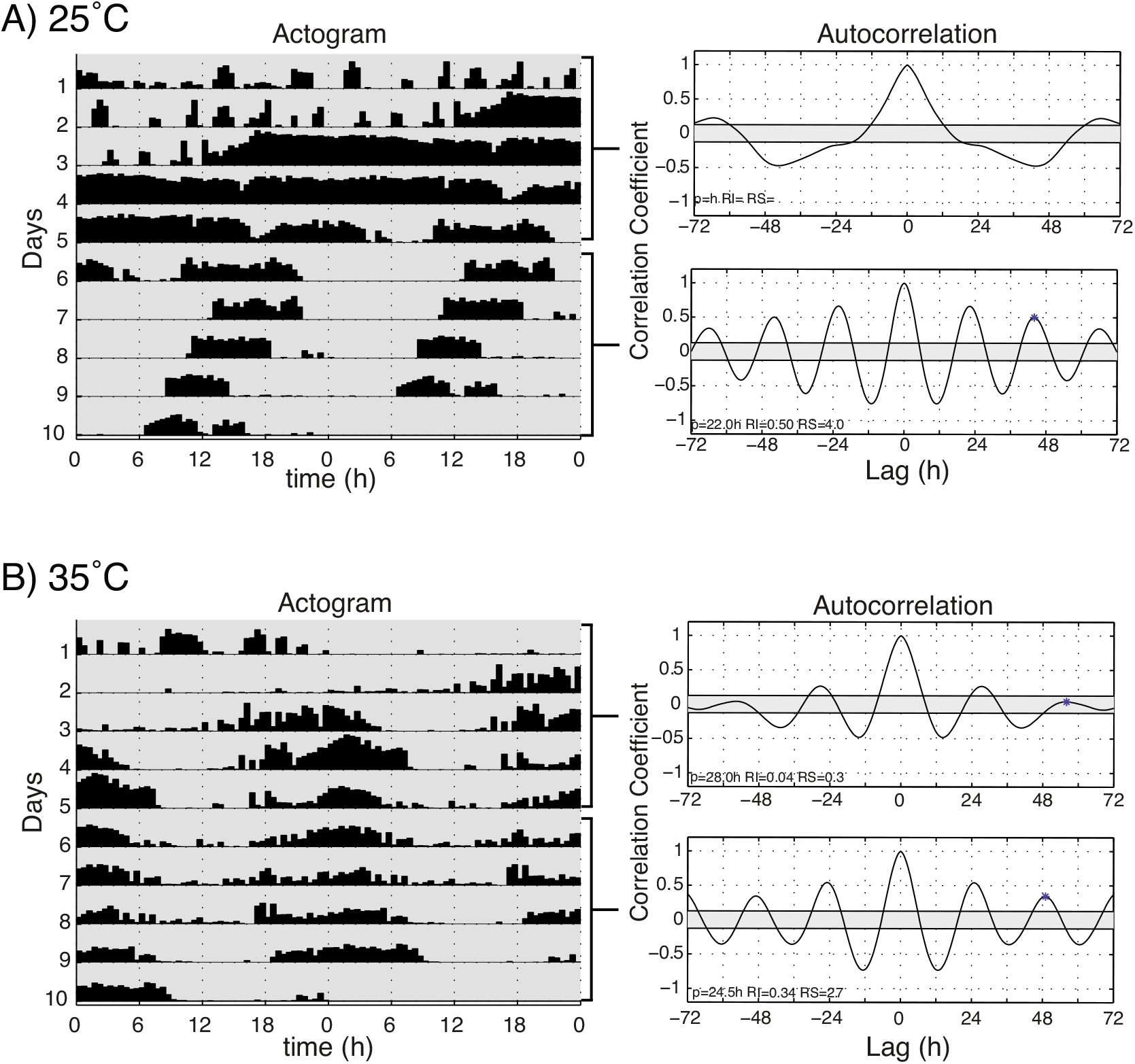
Locomotor activity patterns of young honey bee workers under 25°C or 35°C constant darkness. Double-plotted actograms of representative 1-day-old workers at **A)** 25°C and **B)** 35°C constant darkness. Autocorrelation plots were used to determine rhythmicity of locomotor activity and calculate the endogenous period length (p), rhythm index (RI) and rhythm strength (RS), from days 15 and 6-10 for each individual.

Recent work on different species of honey bees has shown that environmental temperature affects endogenous period length of foragers (Fuchikawa and Isamu Shimizu, 2007; Giannoni-Guzmán et al., 2014). We hypothesized that rhythmic young workers would present endogenous rhythms closer to 24 hours when assayed at 35°C than those assayed at 25°C. To test our hypothesis, we compared the endogenous periods of days 6-10 for rhythmic bees kept at 25°C or 35°C. The resulting analysis revealed that bees kept at 25°C have an average endogenous period length of 23.10hr, compared to that of bees kept at 35°C, whose average period was 24.5hr (Figure 3). This finding is consistent with previous work testing the endogenous period length in foragers (Giannoni-Guzmán et al., 2014; Moore and Rankin, 1985; Spangler, 1972; Toma et al., 2000).

**Figure 3.**
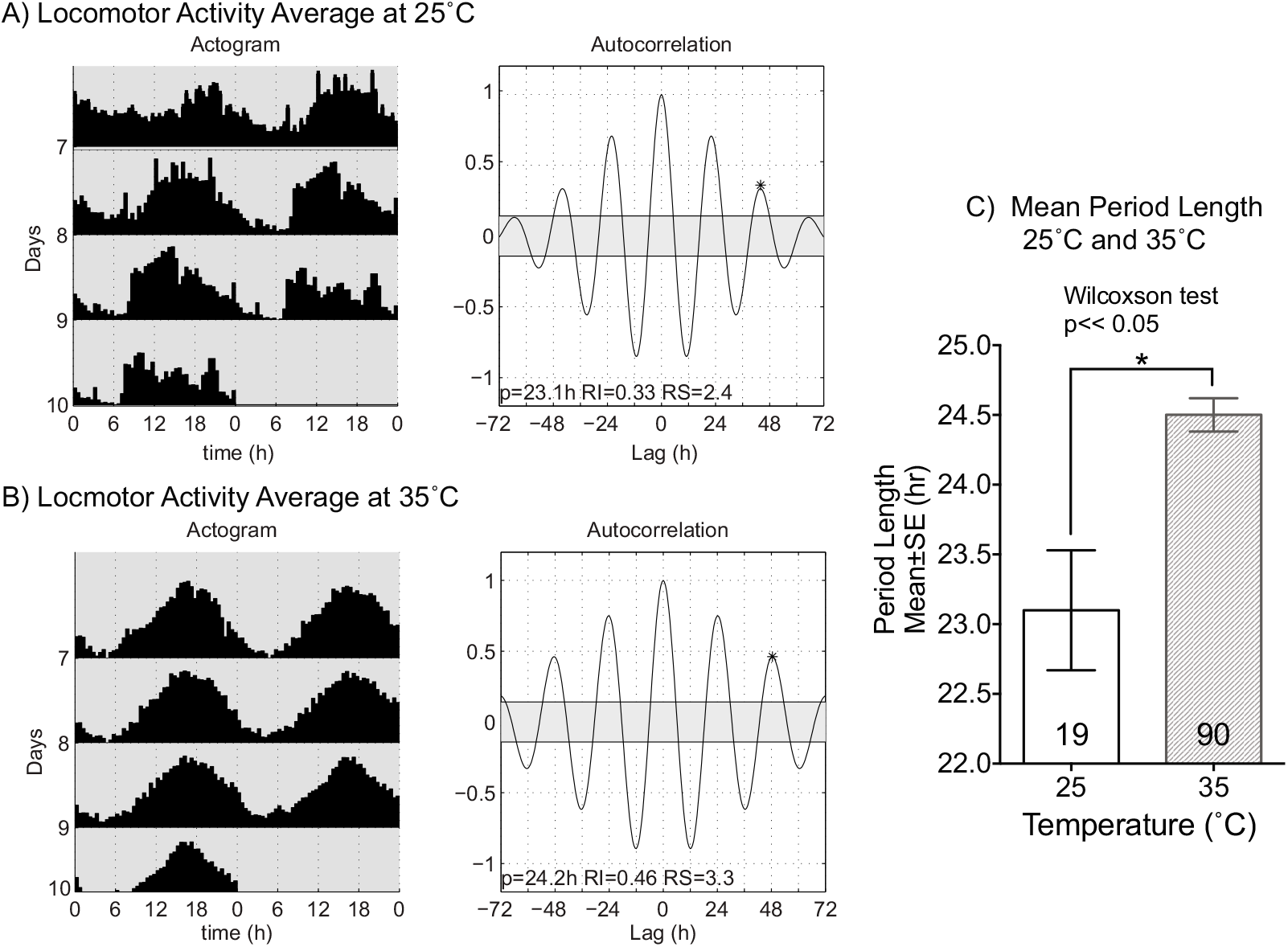
Average endogenous period length of young bees at 35°C is closer to 24 hours. Double-plotted actograms and autocorrelations of the average locomotor profile rhythmic bees at **A)** 25°C and **B)** 35°C for days 6-10. **C**) The mean period length at 35°C (24.5±0.13h SEM) was closer to 24 hours and significantly different from that measured in the 25°C cohort (23.10±0.29h SEM) (ANOVA F=18.59, df=1, p<<0.01).

Interestingly, we observed that the period length standard error of the 25°C group was higher than that of the 35°C group. By observing the distributions of period length for each of the group it was evident that the 25°C group presented a larger degree of variation than the 35°C group (Figure 4A, B). To quantify this variation, we performed Levene’s test of equality of variance, which confirmed that period length in the 25°C cohort varies significantly more than that of the 35°C cohort (F=17.9, df=1, p<<0.01) (Figure 4C). Interestingly, this result does not translate to foragers, where the degree of variation in endogenous period length was not significantly different between foragers at 25°C or 35°C conditions (Levene’s test, F=0.35 df=1, p=0.56) (Figure 4C). Multiple comparisons between young workers and foragers at 25°C and 35°C, revealed that the degree of variation of foragers was similar to that of young workers at 35°C and significantly different from that of young bees at 25°C (Figure 4C). These results suggest that colony temperature after adult emergence plays an important role in the development of circadian circuitry in the honey bee system.

**Figure 4.**
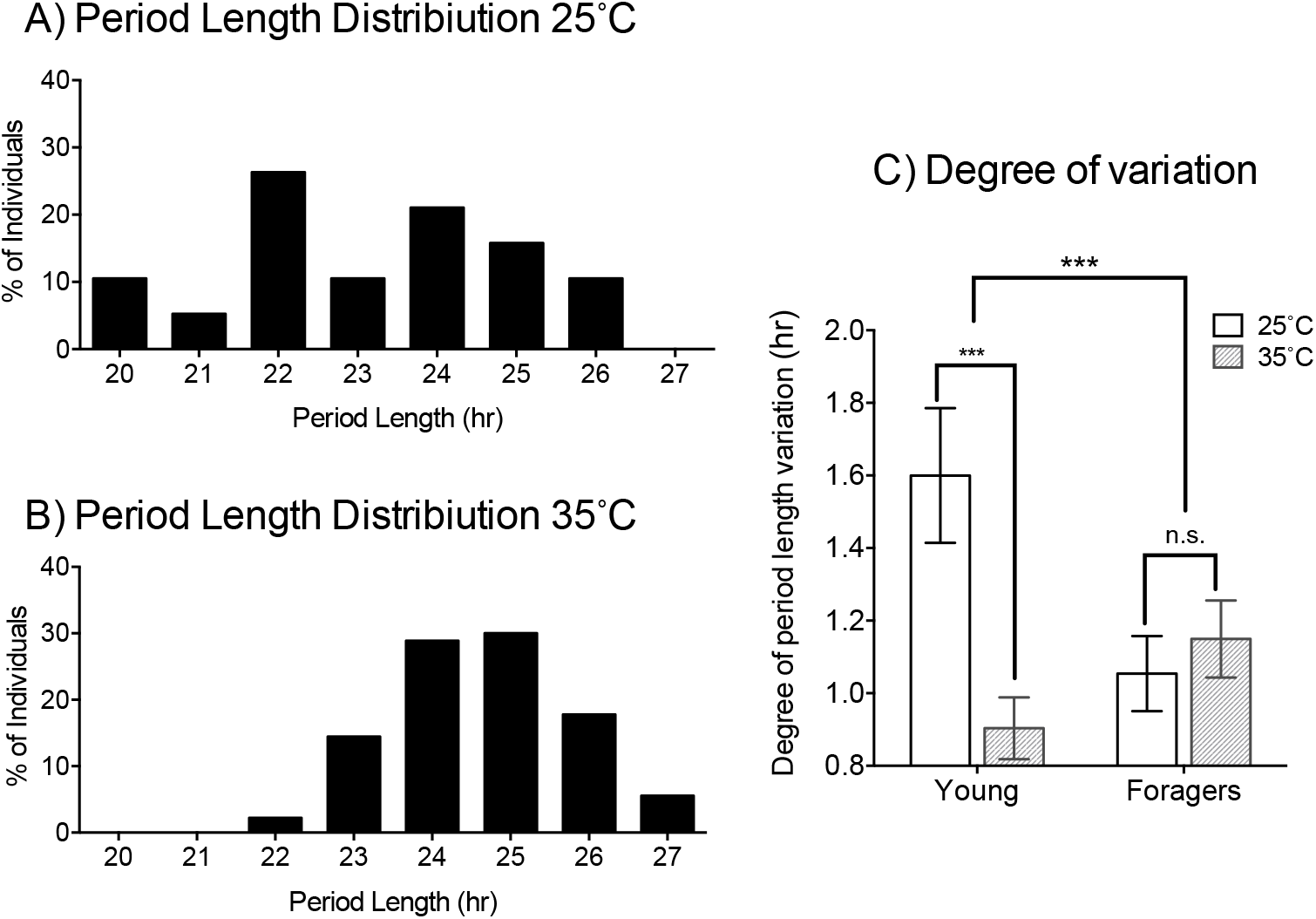
Individual variation of endogenous period length is greater at 25°C than at 35°C. Frequency distributions of endogenous period length of days 6-10 from rhythmic young workers at **A)** 25°C and **B)** 35°C. **C)** Bar graph comparing the degree of period length variation as calculated by Levene’s test of equality of variance for young workers and foragers at 25°C (white bars) and 35°C (gray shaded bars). Significant differences were observed between the young worker cohorts (F=17.9, df=1, p<<0.01), while differences comparison within foragers was not significant (F=0.35 df=1, p=0.56). Multiple comparisons test revealed significant differences (p<0.05) between young workers at 25°C and foragers at either 25°C or 35°C.

During the data analysis of the experiments, another difference that was noticed between the 25°C and 35°C cohorts was their mortality. When we compared the mortality of each group we observed that by day 10 only ~30% of bees in the 25°C cohort survived (Figure 5A). Significantly, this was less than half of the mortality observed in the 35°C cohort, where more than ~65% of the bees were still alive (Gehan-Breslow-Wilcoxon test, p<<0.01). This result is somewhat surprising since our experiments with foragers under the same experimental setup did not reveal significant differences in mortality (unpublished results). Furthermore, by separating each cohort by individuals who developed or did not develop circadian rhythms, we observed a relationship between arrhythmicity and mortality in both groups (Figure 5B). Nonparametric Kruskal-Wallis rank sums test revealed significant differences between arrhythmic and rhythmic individuals at 25° and at 35°C (F=78.13, df=3, p<<0.01). Post hoc analysis using Wilcoxon each pair test uncovered significant differences between 3 of the 4 groups tested, the exception being the comparison of rhythmic individuals at 25°C and arrhythmic individuals at 35°C. In order to ascertain potential factors playing a role in the mortality of honey bee workers, we used a proportional hazards model analysis.

**Figure 5.**
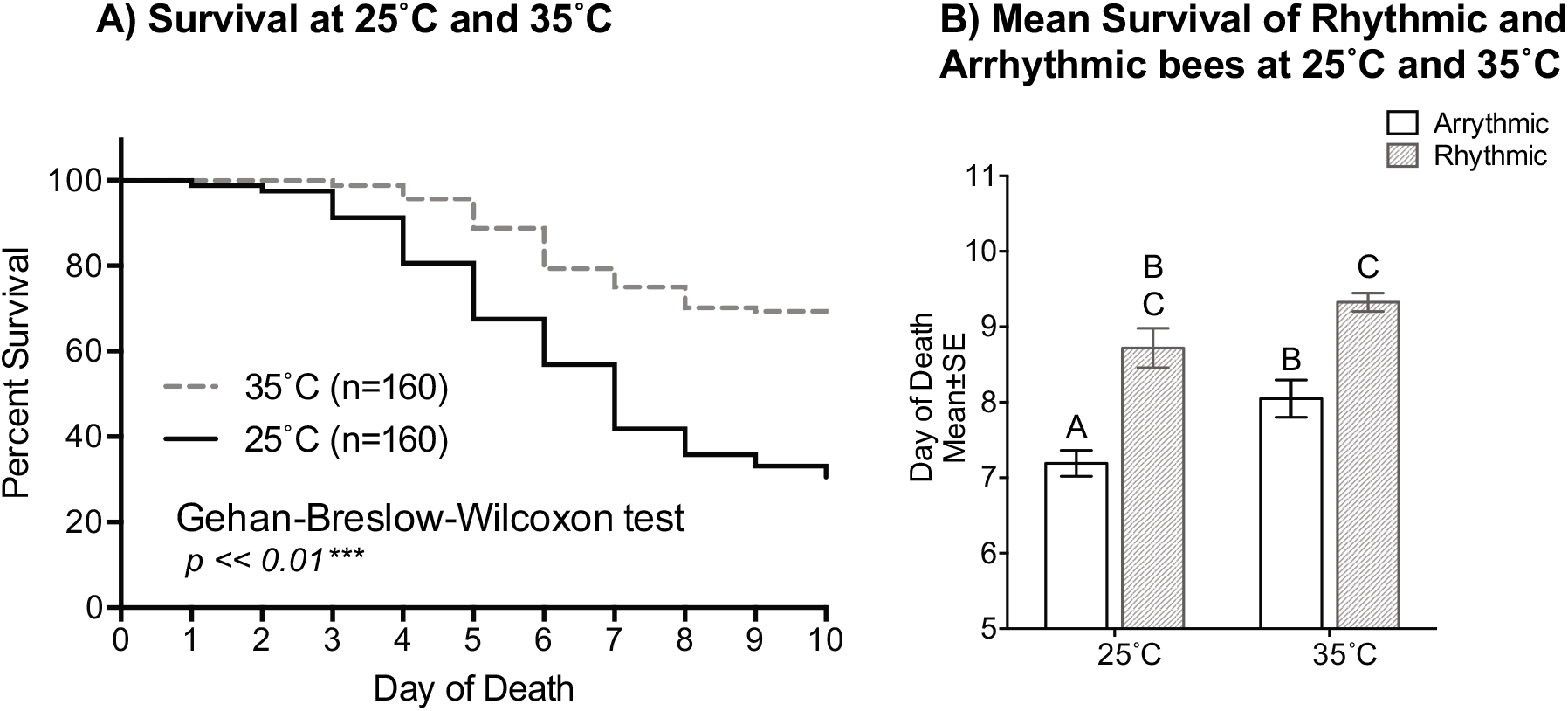
Mortality of isolated young workers greatly increases at 25°C and in arrhythmic individuals. A) Survival plot of 1-day-old honey bee cohorts at 25°C (solid line) and at 35°C (intermittent line). Both visual and statistical comparison of the cohorts revealed that survival of individuals was higher in the 35°C cohort (Gehan-Breslow-Wilcoxon, n=320, p<<0.01). **B)** Bar graph of mean survival and standard error of arrhythmic and rhythmic individuals separated by experimental cohort (25°C or 35°C). Proportional Hazards model revealed that temperature and rhythmicity have independent effects on mortality in young workers (Temperature: X^2^=12.35, df=1, p<<0.001; Rhythm: X^2^=15.64, df=1, p<<0.001; Temperature*Rhythm: X^2^=0.055 df=1, p=0.8142) Wilcoxon each Pair test revealed significant differences (p<0.05) between paired comparisons represented by different letters.

In this analysis environmental temperature, rhythmicity (whether the individual developed rhythms or was arrhythmic throughout the experiment) and the interaction of these factors were tested as the variables causing the observed mortality. The resulting analysis revealed that environmental temperature and rhythmicity, independently, have a significant effect on the mortality of young workers in our assay, while their interaction was not significant (Temperature: X^2^=12.35, df=1, p<<0.001; Rhythm: X^2^=15.64, df=1, p<<0.001; Temperature*Rhythm: X^2^=0.055 df=1, p=0.8142). The combined results suggest that in our experiments mortality is caused by the environmental temperature and the inability to develop a circadian rhythm independently and not their combination.

Our result that temperature positively influences the rate and proportion of individuals developing circadian rhythms combined with the findings from a recent study (Eban-Rothschild et al., 2012) that the first 48 hours in the colony influenced development of strong circadian rhythms led us to postulate the following prediction: If temperature is a key factor in the development of circadian rhythmicity during the first 48 hours after emergence in young workers, then placing 1-day old workers at 35°C for the first 48 hours after emergence and afterwards changing environmental temperature to 25°C, will result in a greater proportion of individuals developing circadian rhythms than 1 day-old workers placed at 25°C. To test this hypothesis, we placed 1-day old bees at either 35°C or 25°C group, after 48 hours, we changed the temperature to 25°C in the first group (35-25°C). Consistent with this prediction we found that exposure to 35°C during the first 48 hours after emergence plays a significant role in the development of circadian rhythms in young workers (Figure 6A). Repeated measures comparison of the cumulative distribution of rhythmic individuals for the 35-25°C group and bees continuously at the 25°C group, was significantly different (F=3.28, df=6, p<0.01). In addition to the effects of temperature on the development of circadian rhythm, we also observed significant differences in the survival of individuals exposed to 35°C for the first 48 hours and those that were kept at 25°C. By day 7 less than 12 individuals had died in the 35-25°C group, while more than 50 had died in the 25°C (Gehan-Breslow-Wilcoxon, n=256, p<<0.01). Taken together, temperature in the colony plays a key role in the development of circadian rhythms of workers.

**Figure 6.**
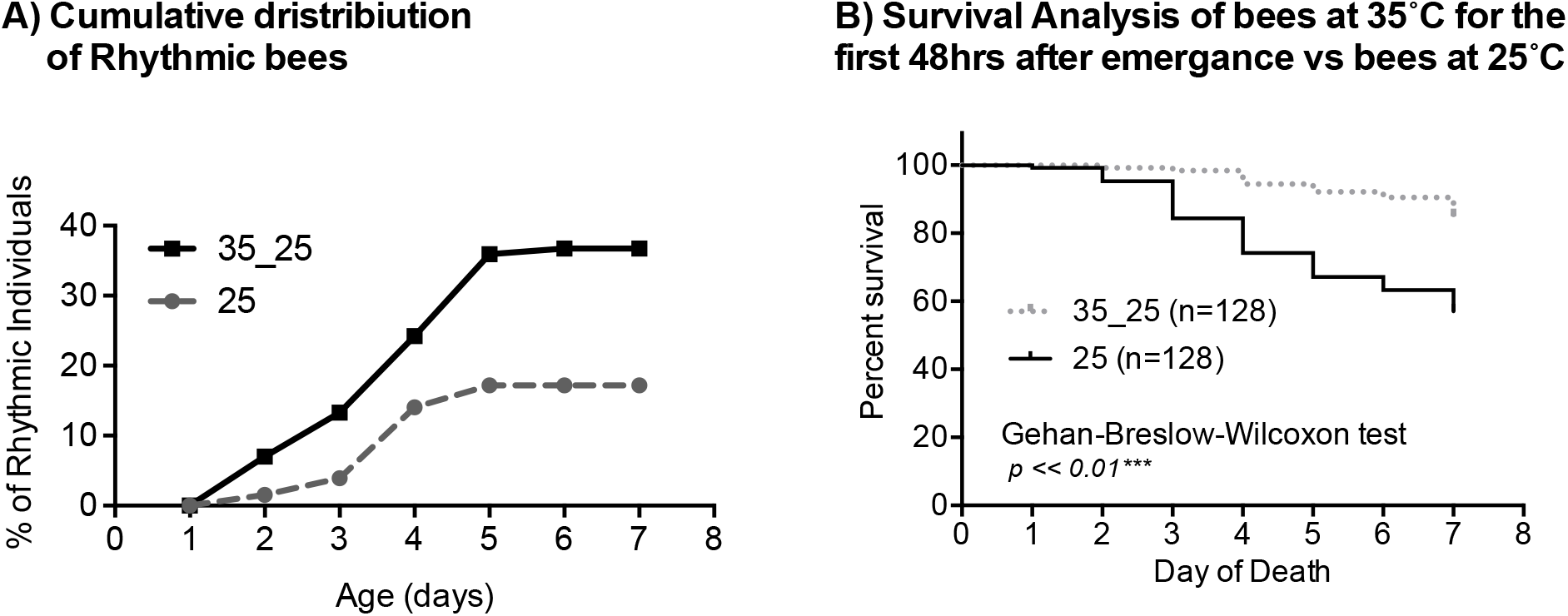
Temperature (35°C) during the first 48 hours after emergence is sufficient to rescue the rhythmicity and mortality effects of 25°C. **A)** Cumulative distribution of rhythmic young workers exposed to 35°C during the first 48 hours after emergence and afterwards placed at 25°C for the remainder of the experiment (35-25°C) compared to that of bees placed at 25°C after emergence. Repeated measures MANOVA revealed significant differences between the rate and proportion of individuals developing rhythmic behavior under these conditions (F=3.28, df=6, p<0.01). **B)** Survival plot of 1-day-old honey bee cohorts at 25°C (solid line) and bees exposed to 35°C for the first 48 hours after emergence (intermittent line). Individuals in the 35-25°C cohort presented significantly better survival rates than bees placed at 25°C since the beginning of the experiment (Gehan-Breslow-Wilcoxon, n=256, p<<0.01).

## Discussion

In the current study, we show that colony temperature plays a key role in the ontogeny of circadian rhythms of young honey bee workers. Previous studies exploring the ontogeny of circadian rhythms of young workers established that circadian rhythms both in the field and in isolation commence around 7-9 days after eclosion (Bloch et al., 2001; Moore et al., 1998; Toma et al., 2000). Experiments that followed uncovered that exposure to the colony environment during the first 48 hours after eclosion significantly impacts the development of circadian rhythms under isolation (Eban-Rothschild et al., 2012). Here we show evidence indicating that the regulation of temperature (~35°C) in the colony is a key social factor determining the development of circadian rhythms. Placing 1-day old workers at the hive’s core temperature in the laboratory results in an accelerated rate and increased proportion of individuals developing rhythmicity (Figure 1). Moreover, we show that this temperature is particularly important during the first 48 hours after eclosion and kept exposure to 35°C in this period is sufficient for early rhythm development (Figure 6). Analysis of the endogenous period length variation of rhythmic bees, suggests that temperature may play a role in the development of the neural circuitry that regulate circadian rhythms (Figures 2,3,4). Lastly, mortality differences between experimental groups were associated with development of circadian rhythms. Taken together, socially regulated temperature plays a key role in the ontogeny of circadian rhythms in honey bee workers.

The proportion and rate of honey bee workers developing circadian rhythms in the 35°C cohort is consistent with work examining the effect of colony environment on circadian rhythms, where after 48 hours of colony exposure between 60-80% of bees presented circadian rhythms (Eban-Rothschild et al., 2012). In addition, individuals exposed to 35°C for the first 48 hours after eclosion presented twice the proportion (40%) of rhythmic bees than bees that at 25°C (20%). This suggests that temperature is one of several factors that play a role in the development of circadian rhythmicity. Furthermore, our results in the 25°C groups are very similar to those of bees that only spent the the first 24 hours after eclosion inside the colony (Eban-Rothschild et al., 2012). Further studies are needed to ascertain the relative importance of temperature on the ontogeny of circadian rhythmicity compared to other colony factors. Studying the neural changes that may occur during the critical period of 24-48 hours after emergence in the honey bee nervous system, may provide clues as to the other factors that influence development of circadian rhythms.

In our data and that of previous studies examining the ontogeny of circadian rhythms in workers, we can observe that not all individuals develop circadian rhythms by the end of the experiment (Eban-Rothschild et al., 2012; Meshi and Bloch, 2007; Toma et al., 2000). While the percent of arrhythmic individuals at 35°C is similar to that of previous studies at the end of the experiment, at 25°C the percent of arrhythmic individuals more than double of that in previous studies (Figure 1). One possible factor that is influencing this result is the time of the year the experiments were carried out, which was winter in Puerto Rico. During winter there are drastic changes in the colony demography and dynamics, such as reduction of brood and complete cease of foraging behavior (Doke et al., 2015). These changes have been mostly studied in temperate zones, where the seasons are marked by drastic changes in weather and may not be necessarily applicable to Puerto Rico. Based on our current data set, further experiments are required to accept or discard the effect of season in the ontogeny of circadian rhythmicity.

Our results and those of other studies provide strong evidence that temperature plays a key role in the ontogeny of circadian rhythms in young workers. However, other studies in the field and laboratory provide evidence that other factors influence the development of circadian rhythmicity, such as genetic background and social environment. Monitoring the behavior of individual bees in the colony as they aged, researchers have shown that rhythmicity, measured using standing behavior as the measure of inactivity in the colony, found that bees of fast genotypes, which show accelerated behavioral development into foragers (Giray and Robinson, 1994), present circadian rhythms as early as 4-7 days of age in some cases, while the slow genotype bees did not show rhythms until the 16-19-day interval (Moore et al., 1998). This finding is consistent with the onset of foraging in slow and fast genotype groups (Giray et al., 1999). The authors of this study conclude that bees inside the hive present rhythmic activity much earlier than onset of foraging. While in the current study we did not control the genetic background of our bees, we did examine two different colonies and obtained similar results.

Since our experiments were performed in the laboratory and individuals were isolated, we cannot measure in the current data set the effects of pheromone on the development of circadian rhythms. However, studies have shown that exposure to the foragers advances development of circadian rhythmicity, while bees housed with young bees of their same age cohort develop rhythmicity later (Meshi and Bloch, 2007). Furthermore, young workers that had direct contact with the brood did not show circadian rhythms even when outside the hive and under light/dark cycles (Shemesh et al., 2007, 2010). Taken together, ontogeny of circadian rhythms in the honey bee colony context is regulated by the socially regulated factors of temperature, social interactions with brood, foragers and young workers, and potentially genetic background.

Endogenous period length of young bees at 35°C (24.5±0.12hr SE) was on average closer to the Earth’s rotational period than that of individuals at 25°C (23.1±0.43hr SE) (Figure 2 and Figure 3). This result is consistent with our previous work on honey bees and work on *Apis cerana* where environmental temperature influenced endogenous period length, suggesting that the circadian clock of young workers is able to compensate for environmental temperature changes (Fuchikawa and I Shimizu, 2007; Giannoni-Guzmán et al., 2014). While differences between 25°C and 35°C cohorts in average period length was consistent with that of foragers, the degree of period length variation was different between young workers at 25°C and 35°C, while the degree of variation in foragers at both 25°C and 35°C was similar to that of young workers at 35°C (Figure 4). Since foraging is the last job a worker performs before dying, these similarities in period length variation between young workers kept at 35°C and foragers at both temperatures is most likely related to foragers having spent the majority of their life inside the colony. The foragers in this study were captured at the entrance of the colony, so we can assume that they had fully developed circadian rhythms. This result suggests that bees exposed to 25°C from a young age may have differences in the development of the circadian network or present a lack of communication between different clocks in the honey bee circadian system.

With regard to the survival rates observed in bees at 35°C and those at 25°C (Figure 5), our analysis indicates that environmental temperature and lack of rhythmicity are independently decreasing survival rates. The effect of temperature on mortality is consistent with our data that temperature is important for the development of circadian rhythmicity and that changes in temperature during development can have long lasting effects later in the honey bee’s life (Becher et al., 2009; Jones et al., 2004; Tautz et al., 2003). It is possible that in addition to development of circadian rhythms, other systems are still under development and do not develop properly at 25°C causing the observed mortality.

Comparing the development of circadian rhythms of honey bee workers with that of other insects suggests that the postembryonic ontogeny in honey bees may be a product of the colony’s social context. Studies examining the circadian rhythms of various insects show rhythmic activity at even pre-adult stages (Fantinou et al., 1998; Kaneko and Hall, 2000; Page and Block, 1980; Tomioka and Chiba, 1982). In the case of crickets and cockroaches, circadian rhythmicity has been documented in pre-adult nymph stages and its patterns change as individuals age (Page and Block, 1980; Tomioka and Chiba, 1982). In other insects such as egg-parasitic wasp *Telenomus busseolae*, adult emergence is timed by their entrainment of light/dark cycles, providing evidence of early development of the circadian system (Fantinou et al., 1998). In contrast to honey bee brood which is kept at almost constant conditions, in these insects the pre-adults (eggs, larvae, pupae, nymphs) are at the mercy of the external environment and having a working circadian system becomes necessary for their survival. In the case of honey bees, since conditions are constant during development, the ability to predict changes in the environment during larval and pupal development becomes less necessary, thus it is possible that honey bee circadian rhythms have evolved to developed after adult emergence when they are needed. For example, in marsupials, such as kangaroos, where gestation is short and many developmental processes occur after birth, the front limbs are much more developed than other systems because upon birth they are required in order to climb to the maternal pouch and to the mother’s nipple to feed (Wittmann, 1981, 1984). At what exact stage of development and what processes are driving the ontogeny of circadian rhythms in honey bee workers is a subject of further research.

In order to present circadian rhythms of locomotion, the connectivity between various systems is necessary. At the brain level, it is known that multiple oscillators that control the timing of locomotor activity, at different times of the day (e.g. morning and evening cells), not only need to communicate but they need to synchronized in a specific manner (Stoleru et al., 2004, 2005). One of the possible processes that may be occurring in the first 48 hours after emergence in workers is the establishment of connections between the multiple oscillators in the honey bee brain. Another circuit that is necessary for locomotor rhythms is the connectivity between motor neurons and the different oscillators in the brain (Blanchardon et al., 2001). Motor neurons are organized forming central pattern generators that coordinate the movement of extremities independently of the brain. However, without a signal from the brain, the initiation and regulation of locomotor rhythms is not possible (Allada et al., 1998). If the formation of this connection is regulated by temperature in honey bees and is occurring in this 48-hour window after emergence, then arrhythmicity may be explained by the failure to establish this connection.

An additional process that is important in the regulation of circadian locomotion is the connections between motor neurons and muscles (i.e. Neuromuscular junction (NMJ). This connectivity has been studied extensively in multiple insects, and the cellular and molecular processes have been well characterized in *Drosophila* (H Keshishian et al., 2003). Experiments exploring the effects of temperature on this connections show a temperature dependent plasticity of motor nerve terminal arborization, where at higher temperatures more arborization of the nerve terminal occurs (Zhong, 2004). In honey bees, measuring circadian gene expression in the brain and muscle of young arrhythmic workers indicates that the muscle clock oscillates, while the brain’s clock did not seem to oscillate (Ben Attia, 2014). Based on what is known in other models and this finding it is possible that different oscillators in the brain have not synchronized with each other and that the connection between the brain’s oscillators and peripheral oscillators, at the time of collection, has not been established and requires further research.

In conclusion, this study shows for the first time the effects of colony temperature on the ontogeny of circadian rhythms, specifically during the first two days after adult emergence. Future studies will examine the weight of temperature as a factor in the development of circadian rhythms and examine the weight of other factors, such as genetic background and social cues. In addition, carefully examining the changes at the neural and gene expression levels occurring during the first 48 hours may provide insight into the mechanisms driving the ontogeny of circadian rhythms in honey bee workers, which remain to be elucidated.

## Acknowledgements

We would like to thank Dr. Arian Avalos and Emmanuel Rivera, for help with the experiments. Thanks Dr. Luis de Jesus for their comments and suggestions. We would also like to recognize the director, Manuel Diaz and the personnel of the Gurabo Experimental Agriculture Station of the University of Puerto Rico at Mayaguez for use of facilities at ‘‘Casa Amarilla’’. This work was sponsored by the National Science Foundation (NSF) awards 1026560, 1633184, 1707355 and the National Institute of Health (NIH) 2R25GM061151-13, P20GM103475.

## Notes

### Competing Interest Statement

The authors have declared no competing interest.

